# Identifying the midline thalamus in humans *in vivo*

**DOI:** 10.1101/2022.02.20.481099

**Authors:** Puck C. Reeders, M. Vanessa Rivera N., Robert P. Vertes, Aaron T. Mattfeld, Timothy A. Allen

## Abstract

The midline thalamus is essential to flexible cognition, memory, and stress regulation and its dysfunction is associated with several neurological and mental health disorders including Alzheimer’s disease, schizophrenia, and depression. Despite the pervasive role of the midline thalamus in cognition and disease, almost nothing is known about it in humans due to the lack of a rigorous methodology for finding this brain region using noninvasive imaging technologies. Here, we introduce a new method for identifying the midline thalamus *in vivo* using probabilistic tractography and *k*-means clustering with diffusion weighted imaging data. This method clusters thalamic voxels based on data-driven cortical and subcortical connectivity profiles, and then segments midline thalamic nuclei based on connectivity profiles established in rodent and macaque tracer studies. Results from two different diffusion weighted imaging sets, including adult data (22-35yrs) from the Human Connectome Project (n=127) and adolescent data (9-14yrs) collected at Florida International University (n=34) showed that this approach reliably classifies midline thalamic clusters. As expected, these clusters were most evident along the dorsal/ventral extent of the third ventricle and primarily connected to the agranular medial prefrontal cortex (e.g., anterior cingulate cortex), nucleus accumbens, and medial temporal lobe regions. The midline thalamic clusters were then bisected based on a human brain atlas (Ding et al., 2016) into a dorsal midline thalamic cluster (paraventricular and paratenial nuclei) and a ventral midline thalamic cluster (rhomboid and reuniens nuclei). This anatomical connectivity-based identification procedure for the midline thalamus offers new opportunities to study this region *in vivo* in healthy populations and in those with psychiatric and neurological disorders.

**Significance Statement:** The midline thalamus is essential for flexible cognition, decision making, and stress regulation. Abnormal development, dysfunction, and neurodegeneration of the midline thalamus is thought to be a critical pathology in several psychiatric and neurological disorders. Yet, little is known about the role of the human midline thalamus. Here we solve the problem of localization *in vivo* by using probabilistic tractography and data driven clustering (k-means), a method capable of capitalizing on known connectivity in non-human animal tracer studies. Localizing the midline thalamus based on its connectivity patterns provides a useful tool for future functional and structural human MRI studies to investigate clinical implications of the midline thalamus in diseases such as Alzheimer’s and schizophrenia.

## Introduction

The midline thalamus is essential to an array of cognitive functions including learning and memory, emotional processing, decision making, and chronic stress regulation (1-4). Dysfunction in the midline thalamus has been directly implicated in numerous neurodegenerative and psychiatric disorders including Alzheimer’s disease (AD), schizophrenia, epilepsy, major depression, and others (5-9). Surprisingly, almost nothing is known about the midline thalamus in the human brain due to the difficulty identifying the location of the midline thalamus in humans *in vivo*.

### Characteristic connectivity of the midline thalamus

The midline thalamus has been extensively studied in rodents and can be separated into four nuclei including the nucleus reuniens (RE), rhomboid nucleus (RH), paraventricular nucleus (PV), and paratenial nucleus (PT) based on histologically determined connectivity and cytoarchitectural features. In both rodents and macaques, the midline thalamic nuclei are characterized by their strong bidirectional connectivity with the agranular mPFC (Fig 1A: Pink), including the infralimbic, prelimbic and anterior cingulate cortex, the nucleus accumbens (nACC; Fig 1A: Purple), and the medial temporal lobe (MTL; Fig 1A: Blue), including the hippocampus, subiculum, parahippocampus, entorhinal cortex, and amygdala. Additionally, the midline thalamus presents little-to-no connectivity with the superior frontal, precentral cortex, and parietal cortex (Fig. 1A) (1, 10-25).

**Figure 1.**
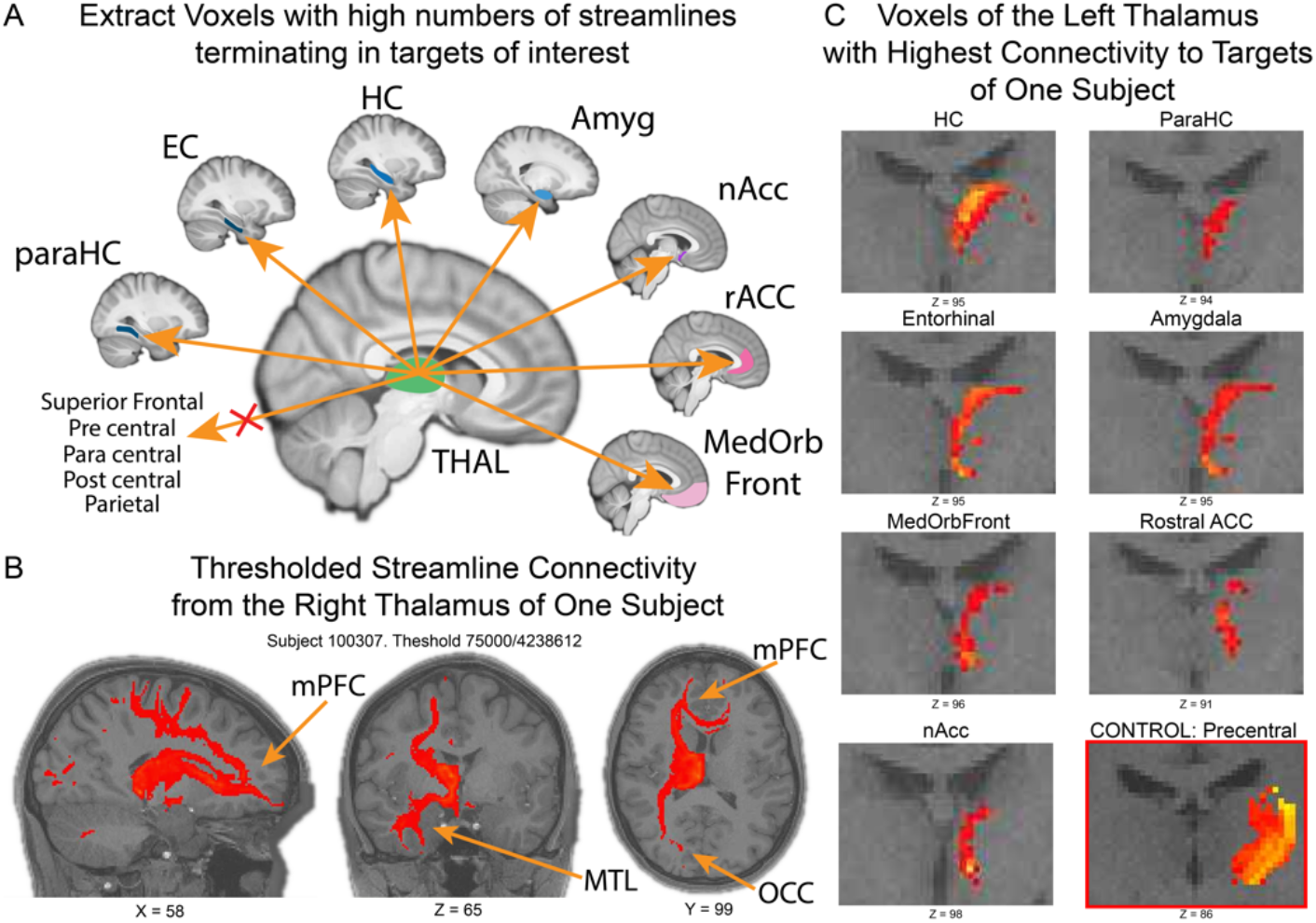
Midline thalamus has high connectivity with mPFC, nACC and MTL, but little connectivity with other regions such as motor and somatosensory cortex (e.g., postcentral and precentral gyrus in the human brain). A) Seed region was the whole thalamus, and target regions of interest (see methods for a list of all regions). B) Streamline density maps showed that the whole thalamus had strong connectivity to MTL regions and mPFC regions and some Occipital region (OCC). Cut off: 75000 from 4238612 total streamlines. C) Thresholding based on visual inspection showed highest connectivity of thalamus to target regions arose from the midline of the thalamus (Thresholding hippocampus 4000/24992 y = 94, parahippocampus 500/6132 y = 94, entorhinal cortex 1700/9721 y = 95, amygdala 3000/24440 y=95, medial orbitofrontal cortex 900/17783 y = 96, rostral anterior cingulate 15/64 y = 91, nucleus accumbens 300/4919 y=98). Connectivity from thalamus to control region revealed connectivity arising from lateral and posterior regions of thalamus (Precentral shown: 8000/19528 y=86).

### Functions of the midline thalamus

The cortical connectivity of the midline thalamic nuclei has driven our understanding of their function. In rodents, studies of the PV and PT often focus on their connections with regions implicated in arousal, stress, and emotional regulation. For example, the PV and PT have been directly implicated in emotional memory (26-28), chronic stress responses (2), and sleep-wake cycles (22; 29). Whereas, RE and RH have been mostly studied in regards to their connections to regions implicated in various learning and memory processes, such as memory consolidation (3), spatial working memory (30, 31), goal directed spatial navigation (32), temporal organization of memory (33), contextual fear memory (34) and behavioral flexibility (31). Interestingly, both PV and RE have neurons that project to both agranular medial prefrontal cortex and the hippocampus, and likely both are related to explicit memory processes under different conditions (35-38).

### Neurological disorders associated with the midline thalamus

Impairments to one or more of the midline thalamic nuclei, or to their connections with cortical and subcortical regions are implicated in numerous neurological diseases including Alzheimer’s disease, chronic stress, epilepsy, and drug addiction. In fact, an early histological study of the human brain in Alzheimer’s disease (6) demonstrated that several midline thalamic nuclei, especially those interconnected with the hippocampus and entorhinal cortex, show dense neurofibrillary tangles early in the course of the disease. This likely impairs communication with the hippocampus and entorhinal cortex, a critical link for memory processing (9) leading to the most severe cognitive deficits (6). That is, the level of pathology found in the midline thalamus in human Alzheimer’s disease patients is correlated with the severity of the noted cognitive impairments, including memory deficits.

### Knowledge gap regarding the human midline thalamus

Currently, there are no rigorous ways to identifying midline thalamic boundaries in humans *in vivo*, making it difficult to study its function in humans. Challenges in the localization of the midline thalamus may be due in part to its small size, location next to ventricles, and lack of clear boundaries in typically obtained magnetic resonance imaging structural scans. Notably, there is no accurate midline thalamic mask available in the field.

Here, we identify the midline thalamus *in vivo* in humans using patterns of anatomical connectivity derived from diffusion weighted imaging (DWI) and probabilistic tractography, informed by known connectivity to regions in the mPFC, MTL and the nACC from macaque and rodent tracer research. The borders of the midline thalamus in humans were delineated utilizing a data driven approach commonly used in machine learning algorithms (k-means clustering). This groundbreaking method provides a midline thalamus mask for researchers to further investigate its function. The results of this approach provide stereotypical midline thalamus coordinates in humans based on highly overlapping connectivity profiles across individual subjects, thus providing some baseline of objectivity for delineation *in vivo*.

## Results

We identified the human midline thalamus by segmenting thalamic voxels based on anatomical connectivity profiles in 127 adults (22-35ys) from the Human Connectome Project with available diffusion-weighted data. First, i*n vivo* connectivity patterns were determined using probabilistic tractography (25,000 streamlines per voxel). This was followed up with a *k*-means clustering analysis that identified voxels with similar connectivity profiles across multiple target regions, as identified in Figure 1A.

### Midline thalamus clusters identified by mPFC, nACC and MTL connectivity profiles

Streamline density maps from representative subjects were produced, i.e., the paths taken from seed (whole thalamus) to target sites to ensure that they followed white matter paths in expected ways and that all potential paths were well sampled (Figure 1B). We further examined the topology of the streamline densities (fdt-map) for several representative participants to assess whether key target regions exhibited robust anatomical connectivity with the entire thalamus (left and right hemispheres analyzed separately). Thresholding of the overall connectivity, based fdt-maps based on visual inspection, showed that the streamlines from the whole thalamus exhibited robust connections with the medial regions of the PFC and regions of the MTL (e.g., see Figure 1B with Subject 100307, right thalamus, thresholded at 75,000 out of a max 4,238,612, Coordinates: x=60, y = 99, z=71, best observed in the coronal and axial slices).

Next, we examined the topography of voxelwise thalamic connectivity to cortical and subcortical targets and control regions (targets: mPFC, nACC, and MTL; controls: superior frontal, pre/post/para central, and parietal cortices) (Figure 1C). We hypothesized that the highest streamlines to the mPFC, nACC, and MTL should arise from the midline thalamic voxels, whereas streamlines to control regions should exhibit preferential connectivity with more lateral and posterior thalamic nuclei voxels. Thresholding based on visual inspection showed that all target regions of interest exhibited robust connectivity to midline thalamic voxels (Figure 1C; Subject 100307; thresholding for the hippocampus 4000/24992 y = 94, parahippocampus 500/6132 y=94, entorhinal cortex 1700/9721 y = 95, amygdala 3000/24440 y=95, medial orbitofrontal cortex 900/17783 y = 96, rostral anterior cingulate 15/64 y = 91, nACC 300/4919 y=98.). In contrast, control regions showed highest connectivity to more lateral and posterior voxels in the thalamus (Fig. 1C; Superior frontal cortex 6500/17223 y=96, pre-central gyrus 8000/19528 y = 86, para central gyrus 900/5860 y = 86, post central gyrus 1200/14837 y = 86, parietal cortex 4000/12421 y = 81).

Although thresholding and selection criteria provided estimates of the location of the midline thalamus, the subjective nature of thresholding combined with variability across subjects did not lend this approach to replicability. Thus, we employed a data driven *k*-means clustering approach, selecting clusters that were defined by preferential connectivity to our key cortical and subcortical regions of interest. This step provided an unbiased, data-driven method for determining the location of the midline thalamus. For each subject and hemisphere, we entered a matrix comprised of cortical/subcortical targets by ipsilateral thalamic voxels into a k-means clustering algorithm set to eight clusters. Across subjects and hemispheres, clusters were consistently observed that were exclusively comprised of midline thalamic voxels (Figure 2A; e.g., the yellow midline cluster in Subj 101006 right image). Next, we selected the specific *k*-means cluster from each participant and hemisphere that matched the target criteria for the midline thalamus. We then binarized and warped the subject specific midline thalamic masks into a template space and averaged the resulting warped masks to calculate a probability of overlap map across our sample. When the resulting mask was thresholded at greater than or equal to 80% overlap results show a striking midline thalamic mask in humans shaped like a ‘reverse C’ bordering the dorsal and ventral extent of the third ventricle (Figure 2B). This approach demonstrates a non-invasive, unbiased method to identify the midline thalamic nuclei across subjects.

**Figure 2.**
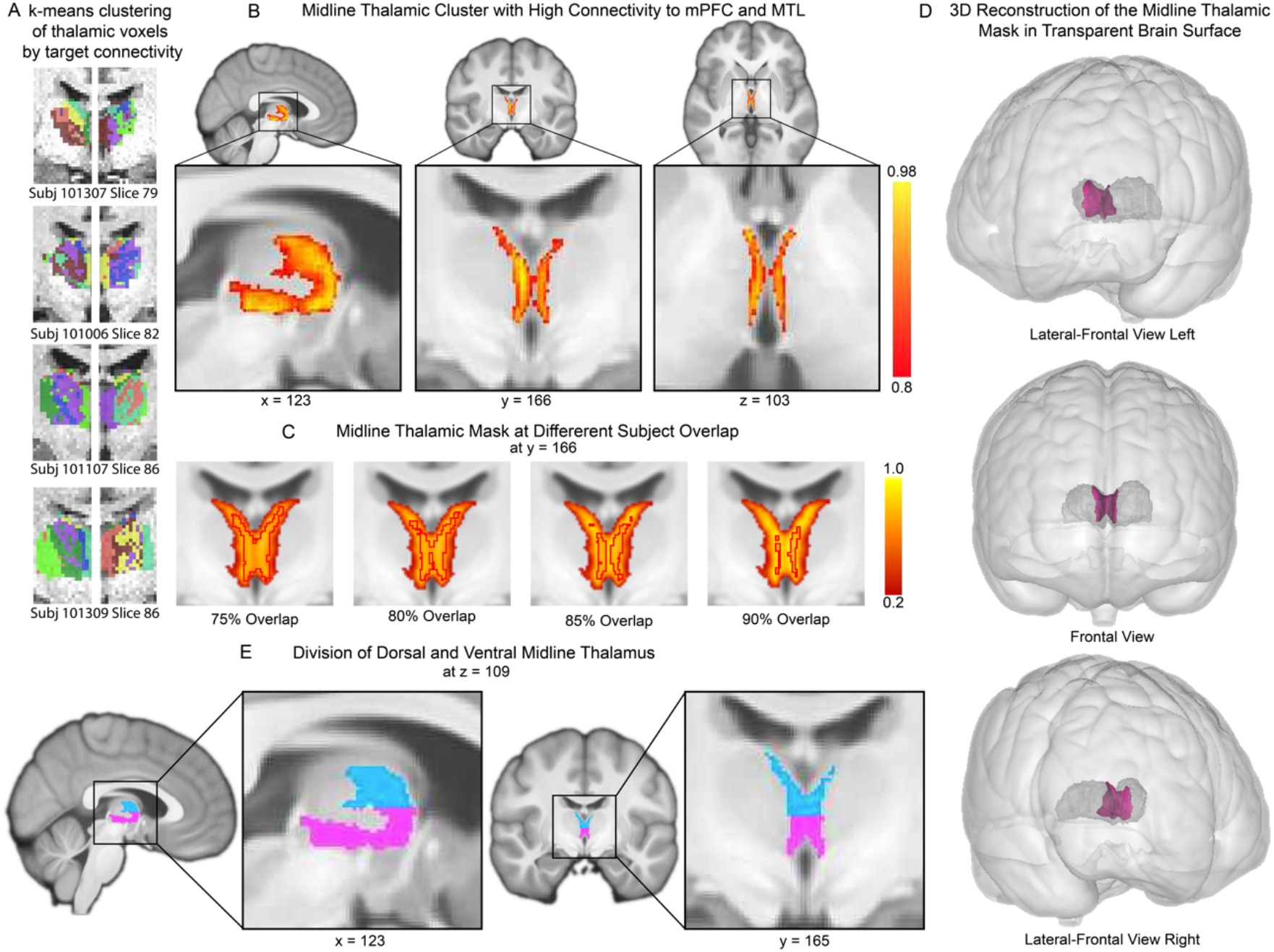
Pinpointing the midline thalamus in humans with *k-*means connectivity clustering. A) Clustering of thalamic voxels with similar connectivity profiles revealed a midline thalamic zone (colors are arbitrary). B) When extracting the cluster with high connectivity to MTL, mPFC and nACC, and not our control regions, we obtained a mask of the midline thalamic mask with over 80% overlap of all participants (80% shown). C) Midline thalamic mask with different overlap in 127 subjects. D) 3D rendering of the midline thalamus (pink) in the thalamus (dark grey) at 80% overlap in 127 subjects. E) Division of the dorsal (blue; PT and PV) and ventral (magenta; RH and RE) thalamus according to the Ding et al., 2016 atlas at z = 109. Where at the point of hemispheric attachment, 48% is dorsal midline thalamus, and 52% is ventral midline thalamus.

### Midline thalamic mask division into dorsal and ventral midline thalamus

Based on anatomical considerations, the dorsal and ventral midline thalamus were split according to an atlas provided by Ding et al. (39). Accordingly, a dorsal/ventral division should be set on the coronal slice where the midline thalamic nuclei paired from both hemispheres. In the Ding et al., atlas (39), the dorsal midline thalamus measured 48% of the entire height of the coronal slice of the midline thalamic nuclei, and the ventral midline thalamus measured 52% of the entire height of the coronal slice of the midline thalamic nuclei. We applied these percentages to divide our midline thalamic mask into a dorsal mask (blue) which measured 52% of the height of the entire midline thalamus on the coronal slice where both midline thalami pair, and a ventral mask (magenta) measuring 48% of the height of the entire midline thalamus on the coronal slice where both midline thalami pair (Fig. 2E).

### Internal replication in adolescent data set

Lastly, we evaluated whether we could extend the identification of the midline thalamus using a locally acquired DWI dataset in 34 adolescents (9-14yrs) (see Methods). Using the same preprocessing and analytic procedures we obtained remarkable reliability in the identification of the midline thalamus (Figure 3). This dataset was smaller compared to the first dataset (N_EMU_=34 versus N_HCP_=127) and with younger participants (M_ageEMU_ = 11 yrs versus M_ageHCP_ = 29 yrs), suggesting that this method can be reliably replicated across dataset of different sizes and age ranges.

**Figure 3.**
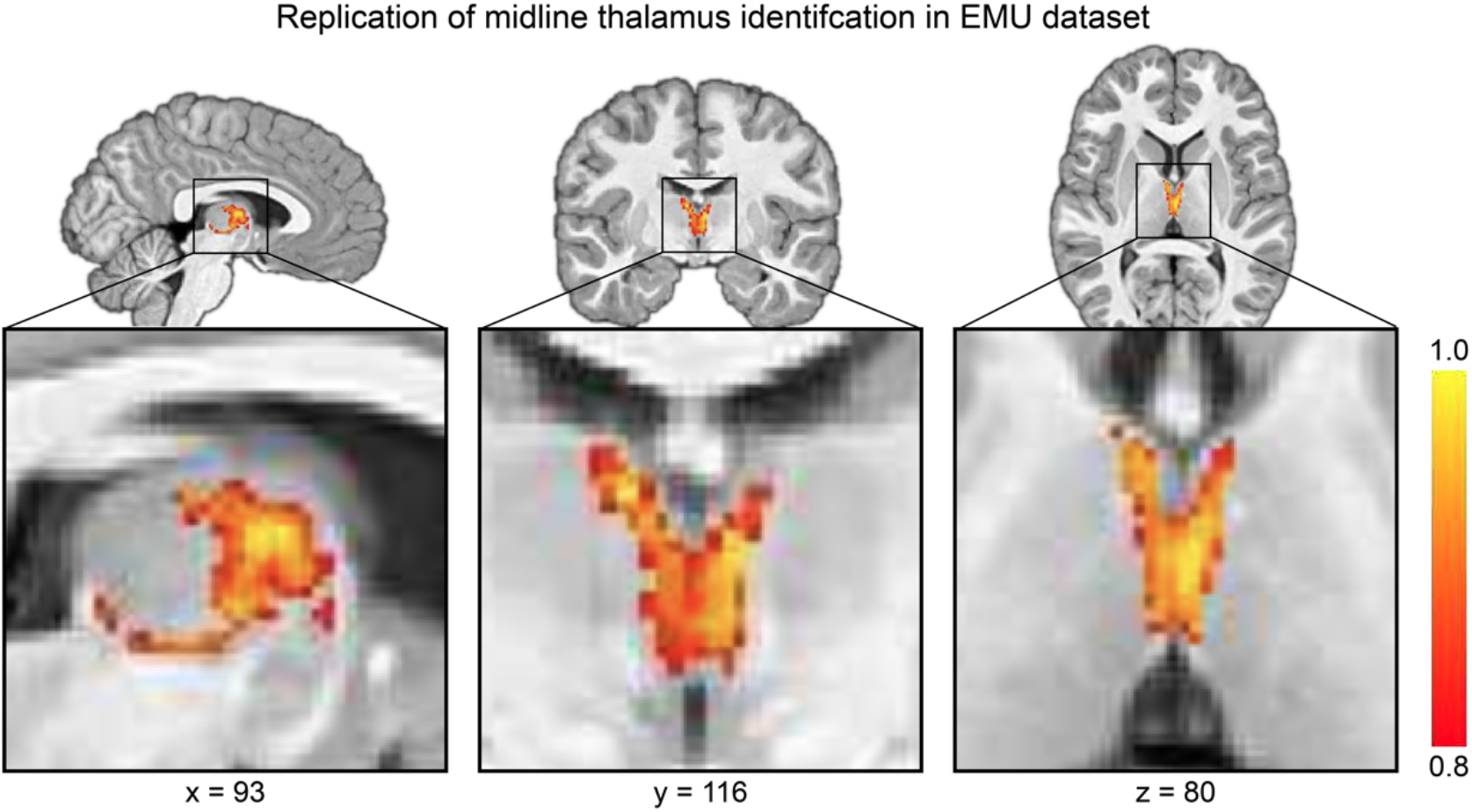
Application and replication of the identification of the midline thalamus to adolescents (9-14yrs) using *k*-means connectivity clustering. Remarkably similar midline thalamic mask of the midline thalamic mask on 34 participants with over 80% overlap of all participants in the dataset (80% shown).

## Discussion

Here, we provide an anatomical identification and segmentation of the midline thalamus in humans *in vivo* based on its known connectivity profile in non-human animals using probabilistic tractography and k-means clustering on diffusion weighted imaging data. The relative location of our midline thalamic mask we found corresponds well with the schematic human atlas by Ding et al., atlas (39) and macaque and rodent tracer studies.

### Clinical applications of the identification of the midline thalamus

This method has obvious clinical applications, as it could help characterize psychiatric and neurological disorders by investigating alterations and changes in the streamline connectivity between the midline thalamus and regions such as the mPFC, HC, or the size of the midline thalamus mask in subjects with dementia and at different stages of Alzheimer’s disease. These measures can in turn be correlated with cognitive and behavioral performances with the use of e.g., memory tasks, learning tasks, mood disorder measures, and cognitive impairment measures or with biomarkers prominent to changes in Alzheimer’s disease.

Additionally, investigation of the midline thalamus during development will provide important insight in neurodevelopmental disorders. Adolescence is a period of increased neural plasticity and hormonal changes that make youth vulnerable to alterations in affect regulation and the development of psychopathologies (40, 41). These cognitive processes arise from coordinated activity between neocortical and subcortical regions (42). Midline thalamic nuclei (e.g., nucleus reuniens) play a role in the coordination of neural activity between the neocortex and medial temporal lobe structures (43). Therefore, identification and segmentation of the midline thalamus *in vivo* can facilitate the investigation of both its structure and function from a neurodevelopmental perspective.

### Future research of the midline thalamus in Alzheimer’s disease

Our study was in part motivated by the absence of an accurate midline thalamic mask for MRI studies that can serve in the investigation of cognitive functions and pathology models. The midline thalamus has shown to be specifically affected by neurofibrillary tangles in Alzheimer’s disease, which presumably impairs communication with HC and EC. These connections are critical for memory processes and disruption in the communication between the midline thalamus and HC and EC could result in memory related cognitive impairments. Our future directions focus on the changes in connectivity between the midline thalamus and the HC at different stages of AD in patients. Additionally, the midline thalamus has been shown to be a critical link closing the communication loop between the mPFC and HC. Specifically, the HC has projections to the mPFC (44), but there are few projections back to HC (reported a recent study; 45). The midline thalamus has strong bidirectional projections with both mPFC and HC (4, 35, 43) and has been shown to be a critical link in this memory dependent circuit (3). It remains unknown whether there are structural impairments between the mPFC and midline thalamus (or mPFC and HC) in Alzheimer’s disease. Using this midline thalamic mask, we can now more accurately study the connectivity patterns in this memory circuit consisting of the mPFC-midline thalamus-HC, both functionally and anatomically in aging, dementia, and Alzheimer’s disease.

### Limitations of the current study

The indirect nature of diffusion weighted imaging used to assess white matter pathways constitutes a limitation of the current study. Tract-tracing and post-mortem histological studies in humans, represent the gold-standard but are limited in efficacy due to ethical and methodological considerations (46). Probabilistic tractography has been previously shown to be robust in dealing with crossing fibers (47) and sensitive to detecting fiber pathways outside of a priori seed based deterministic approaches (48). However, it is impossible to define polarity of the tract, and therefore the direction of fiber pathways (e.g., midline thalamus to HC, vs HC to midline thalamus) is unknown. Lastly, most methods of tractography are sensitive primarily to major pathways and will not always detect small pathways with small inflections. However, Dyrby et al., (49) quantitatively and qualitatively assessed the anatomical validity and reproducibility of in vitro multi-fiber probabilistic tractography against two invasive tracers in the porcine brain. Their findings demonstrated that probabilistic tractography can reliably detected small specific pathways, concluding that DWI probabilistic tractography offers precision in the investigation of structural connectivity (48).

### Replication resulted in remarkable similarity

On a locally collected dataset we applied our connectivity-based identification method and obtained remarkably similar results even though the original and replication datasets were collected from different MRI scanners with different parameters, consisted of different sample sizes and age ranges. The consistency across datasets suggests that the reported identification method is robust and can reliably detect midline thalamic nuclei in a data driven fashion.

### Identifying the midline thalamus is critical to understanding cognitive functions

Demonstrating a robust and replicable method for identifying and segmenting the midline thalamus of the human brain is necessary. It can not only assist in the functional investigation of midline thalamus in cognition, including memory processes and chronic stress regulation and arousal, but also in the structural integrity of neurological disorders known to exhibit midline thalamic pathology, such as Alzheimer’s disease.

## Materials and Methods

### Participants & Image Acquisition

#### Acquisition parameters HCP dataset

The WU-Minn dataset from the Human Connectome Project (HCP; 50) were obtained through “connectome in a box” (www.store.humanconnectome.org). We selected the first 127 participants from the HCP dataset. Ages ranged from 22 to 35 years old (M = 29.000, SD = 3.432) from which 72 were female. Participants were scanned on a customized Siemens 3T “Connectome Skyra” at Washingtoon University in St. Louis, using a 32-channel Siemens head coil. The structural image had an acquisition time of 7:40 min (repetition time; TR = 2400 ms, echo time; TE = 2.14 ms, inversion time, TI = 1000 ms, flip angle = 8°, FOV = 224×224 mm, voxel size = 0.7 mm isotropic, BW of 210 Hz/Px). The diffusion weighted images included 6 runs of approximately 9 minutes and 50 seconds. There were 3 different gradient tables. Each table was acquired once with right-to-left and left-to-right phase encoding polarities, respectively. Each table has approximately 90 diffusion weighting directions plus 6 b=0 acquisitions interspersed throughout each run. Diffusion weighting consisted of 3 shells with b = 1000, 2000 and 3000 s/mm^2^ interspersed. The sequence was a spin-echo EPI (TR = 5520 ms, TE = 89.5 ms, flip angle = 78°, FOV = 210×180 mm, matrix = 168×144 mm, and refocusing flip angle of 160 degrees). There were 111 slices and were 1.25 mm thick with 1.25 mm isotropic voxels. The multiband factor was 3 and 0.78 ms of echo spacing was used. Phase partial Fourier was 6/8.

#### A*cquisition parameters EMU dataset*

This dataset was obtained from the EMU project at FIU and contained data from 34 children between 9 to 14 years of age (M = 11.4 SD = 2.0) from which 16 were female. Data was collected on a 3T Siemens MAGNETOM Prisma scanner using a 32-channel head coil at the Center for Imaging Science (CIS) at Florida International University (FIU). The T1-weighted (T1w) data was collected using a magnetization-prepared rapid gradient echo sequence (MPRAGE: TR=2500ms, TE=2.9ms, flip angle=8º, FOV=256mm, 176 sagittal slices, voxel size=1mm isotropic). The diffusion weighted data acquisition was 1.7 mm isotropic and a multiband EPI was acquired. Slice acceleration factor = 3, 96 directions, seven b=0 frames, and four b-values: 6 directions with 500s/mm^2^, 15 directions with 1000s/mm^2^, 15 directions with 2000s/mm^2^ and 60 directions with 3000s/mm^2^. Additionally, a field map opposite of the phase encoding direction of the DWI acquisition was acquired for the correction of b=0 distortion.

### Data Processing and Analysis

#### Preprocessing HCP dataset

HCP data was preprocessed by the HCP. Using the structural images, they produced an undistorted “native” structural volume space for each participant, performed a bias field (B1) correction, and lastly registered the native structural volume to Montreal Neurological Institute (MNI) space. FreeSurfer (5.3.0-HCP) was used to segment the volume into predefined structures, reconstruct white and pial cortical surfaces, and perform standard folding based surface registration to the surface atlas (fsaverage). Their diffusion preprocessing included normalization of the b0 image intensity across six diffusion series using the “topup” tool (51), removing EPI distortions. FSL’s EDDY algorithm was used for eddy-current-induced correction and motion correction. Next, they did gradient nonlinearity correction and removed spatial distortion and mean b0 image was corrected from distortion. Bvalue and bvector deviation were calculated. They registered the mean b0 to native volume T1w with FLIRT and bbregister and transformation of diffusion data, gradient deviation, and gradient directions to 1.25 mm structural space. Brain mask was based on the segmentation output from FreeSurfer.

#### Preprocessing EMU dataset

Each participant’s T1-weighted structural scan was skull-stripped and registered to an MNI-152 template via a rigid body transformation. This was used to reduce normalization errors due to large differences in position across participants and to generate a template close to a commonly used reference. The rigid-body transformed structural scans were then used to create a study-specific template using ANTs (53). Following template generation, each participant’s skull-stripped brain in FreeSurfer space was normalized to the study template using ANTs non-linear symmetric diffeomorphic mapping. The warps obtained from normalization were used to move binarized midline thalamic masks following k-means clustering into the study-specific template space for group-level averaging.

For diffusion image preprocessing, each participant’s diffusion weighted scan was registered to FreeSurfer structural space using boundary-based registration with the reference image being the first acquired b=0 frame. The FreeSurfer parcellation and segmentation file (aparc+aseg) was then binarized and transformed into diffusion EPI space using FreeSurfer’s ApplyVolTransform tool and was then binarized and dilated by 1mm to include edge voxels to act as a brain mask. Susceptibility distortion correction was then performed using FSL Topup, followed by eddy current correction using FSL Eddy on the diffusion data masked by the diffusion EPI space brain mask.

#### Probabilistic Tractography analysis of both datasets

Next, we used Bayesian Estimation of Diffusion Parameters (BEDPOSTX; 54) on the preprocessed DWI data to model crossing fibers within each brain voxel and to create a fitting of the probabilistic diffusion model on corrected data, separately, on both datasets. The results of BEDPOSTX were the basis of all subsequent probabilistic tractography based analyses. The thalamus of both hemispheres was separately used as seed regions for probabilistic tractography with ipsilateral cortical and subcortical targets obtained from Freesurfer. The ipsilateral targets were precentral gyrus, postcentral gyrus, lateral frontal cortex (caudal middle frontal, superior frontal, rostral middle frontal, lateral orbitofrontal, pars orbitales, pars triangularis, frontal pole), parietal cortex (supramarginal, superior parietal, inferior parietal), temporal cortex (inferior temporal, middle temporal, superior temporal, temporal pole, insula, transverse temporal), occipital cortex, paracentral gyrus, caudal anterior cingulate gyrus, medial orbitofrontal, rostral anterior cingulate gyrus, superior frontal gyrus, medial posterior cortex (posterior cingulate, precuneus, isthmus cingulate), medial occipital cortex (cuneus, lingual, pericalcarine), fusiform gyrus, entorhinal cortex, parahippocampal gyrus, hippocampus, amygdala, and nucleus accumbens). We used ProbtrackX to run probabilistic tractography on the data. Briefly, ProbtrackX produces sample streamlines, starting from a voxel within the seed region, and then iterates between 1) drawing an orientation from the voxel-wise bedpost distributions and 2) taking a step in the direction of diffusion orientation and 3) checking for any termination criteria (see below). These sample streamlines will then be used to investigate how often a streamline from each voxel of the thalamus ends up in one of the target regions. This streamline distribution can be thought of as an estimated connectivity distribution.

We sampled 25000 voxel-wise principal diffusion directions (streamlines) from each voxel in the seed region (the whole thalamus), each time computing a streamline through these local samples to generate a *probabilistic streamline* from the distribution on the location of the true streamline. The termination criteria where either when a streamline hit a voxel in the target mask, or 2) reaching 5000 steps; and restriction criteria were that the streamline random walks could not go backward to the previous voxel and were restricted in how sharply pathways can turn in order to exclude implausible pathways (minimum angle of 90 degrees). FMRIB’s Diffusion Toolbox (FDT) was then used to create a *connectivity distribution*. Primarily, we recorded the target destination of each random walk for every thalamic voxel. We used anatomical derived from Freesurfer anatomical regions of interest (surfer.nmr.mgh.harvard.edu) using their aparc+aseg files. Cortical and subcortical target files were binarized, transformed using a nearest neighbor interpolation, and resampled to DWI resolution prior to running the ProbtrackX algorithm. Masks of the ventricles were used as avoid masks. Next, we made a voxel by target matrix with the frequency of how often each voxel in the thalamus made it to each specific ipsilateral target which were entered into the k-means clustering algorithm.

#### K-means clustering on both datasets

The voxel by target (a.k.a., feature) multidimensional data was entered into a *k*-means clustering algorithm using scikit-learn (https://scikit-learn.org/stable/), which clusters data by separating samples into n groups of equal variances, minimizing inertia (within-cluster sum-of-squares) we constrained the solution to 8 clusters. Voxels with similar target patterns of connectivity based on how many times their random walks made it to the set of targets were grouped together into a k-means cluster. K-means clusters with connectivity profiles that preferentially included high connectivity to targets of interest and minimal connectivity with targets of non-interest were used to select k-means clusters that matched the connectivity profile informed by the non-human animal studies. Targets of interest were medial temporal lobe regions (hippocampus, parahippocampus, entorhinal cortex, and amygdala), nucleus accumbens and medial prefrontal cortex regions (medial orbitofrontal cortex, rostral anterior cingulate cortex; Fig. 1A). These mPFC labels correspond to agranular cortex Brodmann areas 14 (orbital prefrontal cortex), 25 (infralimbic cortex), 32 (prelimbic cortex) and 24 (anterior cingulate gyrus) according to Passingham and Wise (23). Targets of non-interest included superior frontal cortex, precentral gyrus, paracentral gyrus, post central gyrus, parietal cortex (Fig. 1A). The resulting k-means clusters that met our target/non-target criteria were binarized and normalized to the MNI template, averaged across our sample, and thresholded to select voxels that only exhibited 80% or greater overlap across all participants. We divided the resulting thresholded midline thalamus mask based on Ding et al., (39) atlas into a dorsal midline thalamus mask (upper 48% of pairing height in the most anterior slice where the midline thalamus pairs) and ventral midline thalamus mask (lower 52% of the pairing height of the most anterior slice where the midline thalamus pairs). The dorsal midline thalamus mask includes the PT and PV, and the ventral midline thalamus mask includes the RH and RE. This results in three masks per hemisphere that can be used for further analyses including fMRI or structural connectivity analyses: a full midline thalamus mask, a dorsal midline thalamus mask, and a ventral midline thalamus mask.

## Acknowledgments

We thank Adam Kimbler and Nathan Muncy for help with the analytical scripts.

